# *Lingyuanfructus*: The First Gymno-angiosperm

**DOI:** 10.1101/2022.05.27.493677

**Authors:** Xin Wang

## Abstract

Distinct from gymnosperms with naked ovules, angiosperms are defined and characterized by their enclosed ovules. According to Darwinism, angiosperms should be derived from their ancestors that have exposed ovules. Theoretically and precisely, such a plant is expected to have started but not yet completed enclosing its ovules. This expectation is hitherto never met by fossil evidence. Here I report a fossil plant, *Lingyuanfructus hibrida* gen. et sp. nov, from the Yixian Formation (Lower Cretaceous) of Liaoning, China meeting this expectation. With ovules naked and enclosed in a single specimen, *Lingyuanfructus* blurs the former distinct boundary between angiosperms and gymnosperms.

## Introduction

The origin of angiosperms has been a focus of botanical debates ^1-33^ because it is closely hinged with a robust natural angiosperm systematics, which covers more than 300,000 species (more than 90% species diversity of land plants). Theoretically, angiosperms should be derived from their non-angiospermous ancestors with naked ovules, namely, a plant supposedly transitional between angiosperms and gymnosperms should have started but not yet completed its ovule-enclosing, which means that both enclosed and naked ovules occur in a single plant. So far, such an expected evolution snapshot exists only theoretically. The Yixian Formation (Barremian-Aptian, Lower Cretaceous) has yielded diverse angiosperms ^1-5,8,17,24,28,30,31,34-37^ and thus is a promising bonanza for solving the problem of origin of angiosperms. Here I report a new angiosperm, *Lingyuanfructus hibrida* gen. et sp. nov, from the Yixian Formation of Lingyuan, Liaoning, China. *Lingyuanfructus* has both exposed and enclosed ovules in a single specimen, just as expected above. Although its final affinity may be controversial for a while, *Lingyuanfructus* does blur the formerly distinct boundary between gymnosperms and angiosperms.

## Materials and Methods

The specimen was collected from the Yixian Formation near Dawangzhangzi Village, Lingyuan, Liaoning, China (41º15 ꞌN, 119º15 ꞌE), including a specimen of two facing parts of gray siltstone preserved as a compression with little coalified residue. Previously, *Archaefructus sinensis*^3^, *Sinocarpus decussatus* ^4,5^, *Nothodichocarpum lingyuanensis*^30^, *Neofructus lingyuanensis*^17^, and *Callianthus dilae* ^24^ have been uncovered from the locality. The specimen was photographed using a Nikon D200 digital camera, and a Nikon SMZ1500 stereomicroscope with a Nikon DS-Fi1 digital camera. SEM was performed using a MAIA3 TESCAN housed at the Nanjing Institute of Geology and Palaeontology, Nanjing, China. All figures were organized using a Photoshop 7.0.

## Results

*Lingyuanfructus* gen. nov

### Diagnosis

Distal portion of plant including branch, leaves and carpels. Branch slender and straight. Leaf strap-shaped, smooth-margined, parallel-veined, with rare mesh. Flower pistillate, including paired carpels. Ovules multiple per carpel, enclosed, rarely exposed.

*Lingyuanfructus hibrida* gen. et sp. nov (Figures 1-3)

**Figure 1.**
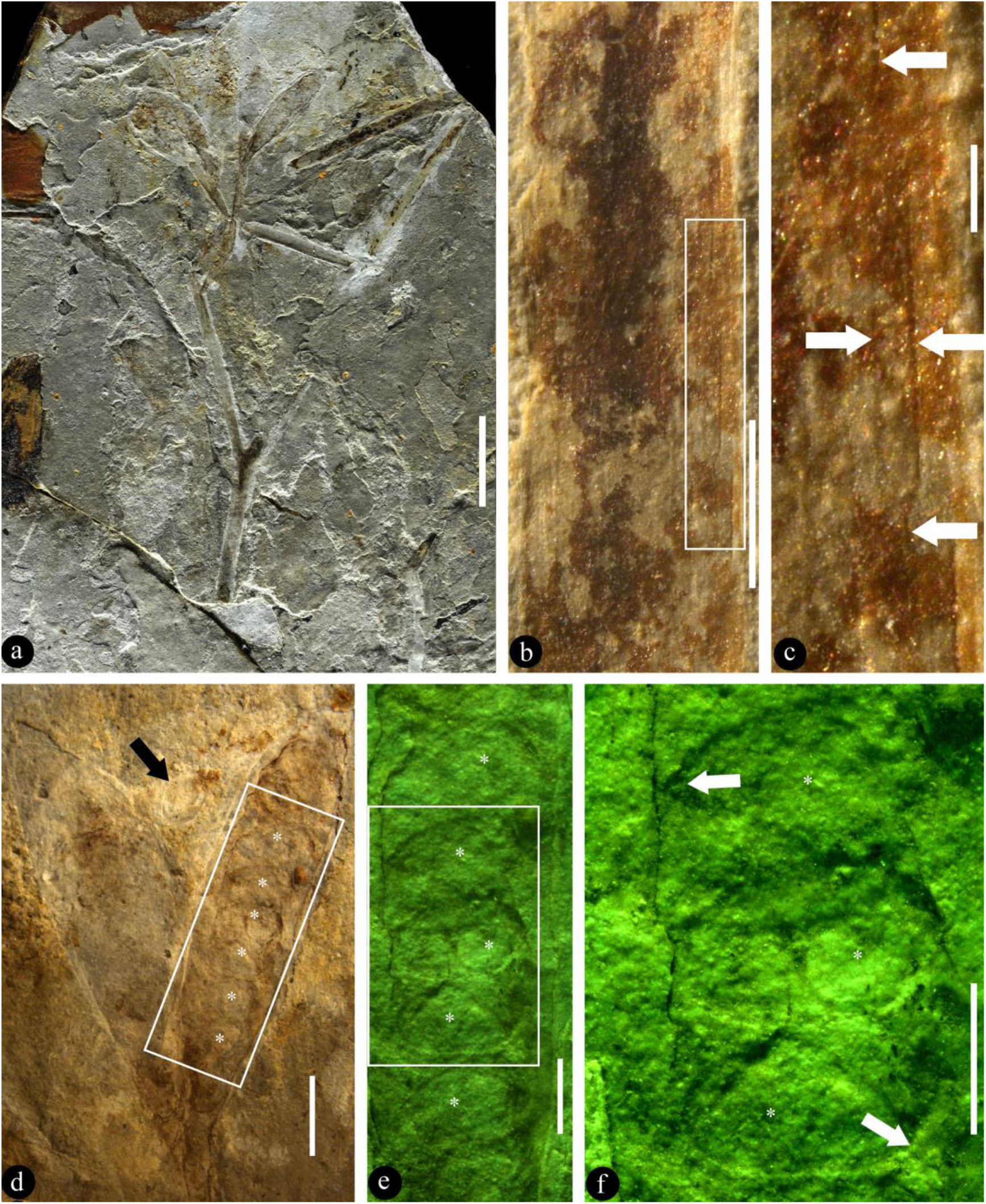
*Lingyuanfructus hibrida* and its details. B-F, stereomicroscopy. **A**. Holotype of *Lingyuanfructus*, including physically connected branch, leaves, and carpels. Refer to Fig. 3a. PB23898a. Scale bar = 1 cm. **B**. Strap-shaped leaf with smooth margins and parallel venation. Scale bar = 1 mm. **C**. Parallel venation and a rare mesh (arrows), enlarged from the rectangle in Fig. 1b. Scale bar = 0.2 mm. **D**. Paired carpels with an ovule (arrow) outside and ovules (asterisks) inside the right carpel. PB23898a. Scale bar = 2 mm. **E**. Detailed view of the rectangle in Fig. 1d, showing ovules (asterisks) inside the carpel. Scale bar = 1 mm. **F**. Multiple ovules (asterisks), with opposite orientations (arrows), inside a single carpel. Scale bar = 1 mm.

### Diagnosis

The same as the genus.

### Description

The fossil is a compression, including two facing parts, embedded in yellowish siltstone. The fossil is 8.3 cm long, 3.7 cm wide, including a branch, leaves, and at least two pairs of carpels physically connected (Fig. 1a). The branch includes at least three internodes, up to 1.7 mm in diameter basally, weakly tapering to the distal (Fig. 1a). An internode is up to 21 mm long and 1.5 mm wide (Fig. 1a). The leaves are strap-shaped, up to 20 (44?) mm long and 1.3 mm wide, smooth-margined, parallel-veined, with rare mesh (Figs. 1a-c). At the branch terminal, there is a 7 mm long shared stalk that furcates into two approximately 7.5 mm long separate stalks (Fig. 1a). Each of the separate stalks bears a pair of carpels, smoothly transitional to the carpels, showing no trace or scars of perianth or other lateral appendages (Figs. 1a, d). Each carpel is 9-11 mm long and 2.8-3.7 mm wide (Figs. 1a, d-e, 2a). There are 10-15 ovules in each carpel, attached to both margins of the carpel (Figs. 1a, d-f). Each ovule is oval-shaped, 0.6-2.5 mm long and 0.6-1.4 mm wide (Figs. 1d-f, 2a). Although most ovules are enclosed in the carpels, two of the ovules are exposed and attached to the adaxial margin of the carpel (Figs. 1a, d, 2a, 3a). These two ovules are 2-2.3 mm long, 1.26-1.5 mm in diameter, orthotropous, unitegmic, without obvious funiculus (Figs. 1d, 2a-d). The integument is 0.34 mm thick and 1.2 mm long, while the nucellus is 0.93 mm long and 0.65 mm in diameter, and free from the integument except at the base (Figs. 2b-d). The micropyle is approximately 0.5 mm wide (Figs. 2b-d).

**Figure 2.**
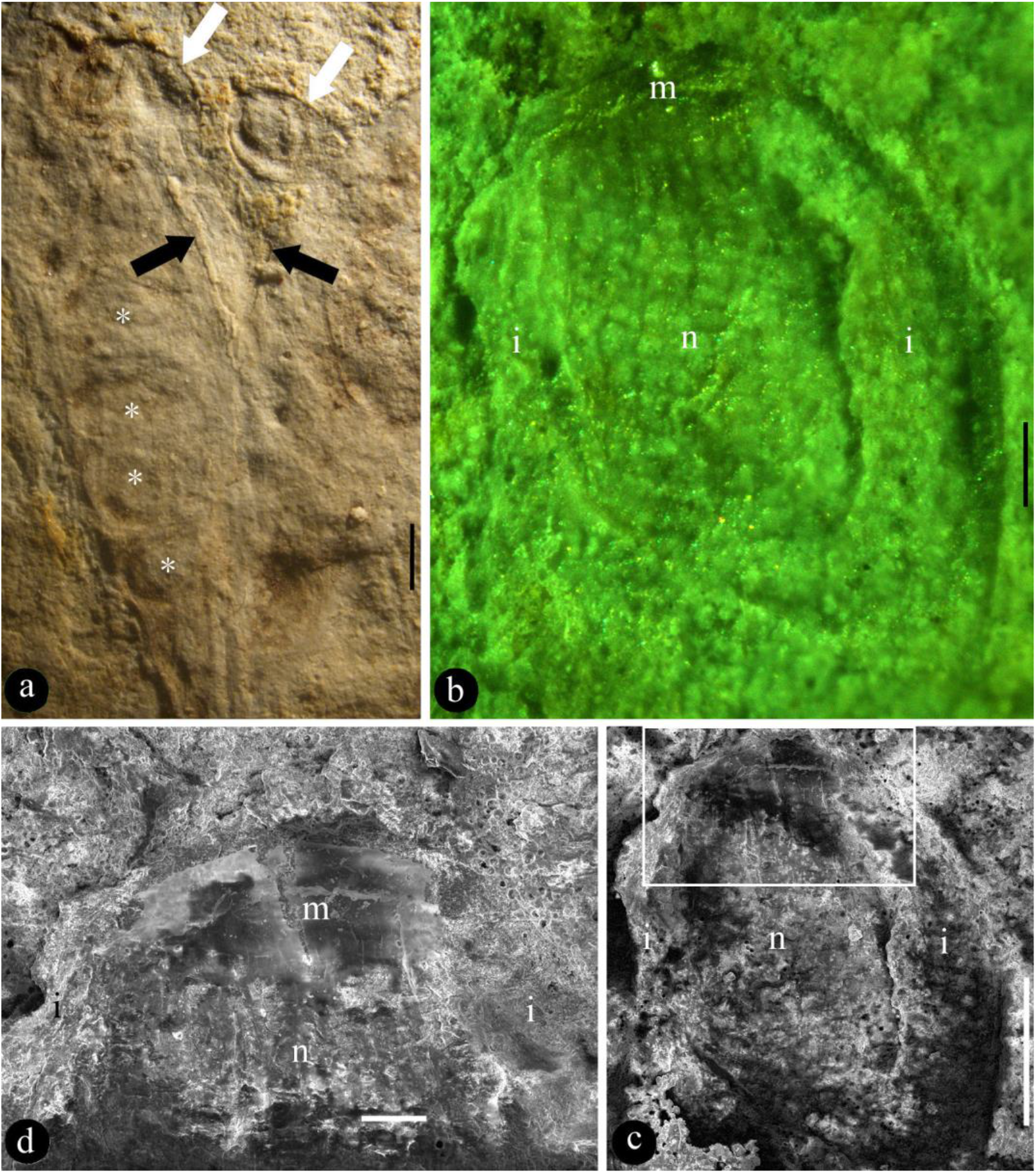
Details of the exposed ovules of *Lingyuanfructus hibrida*. A-B, stereomicroscopy; C-D, SEM. **A**. Two exposed ovules (white arrows) connected to adaxial carpel margins (black arrows). Note ovules (asterisks) enclosed in a carpel. PB23898b. Scale bar = 1 mm. **B**. The orthotropous unitegmic ovule outside carpel shown in Fig. 1d, showing nucellus (n), integument (i), and apical micropyle (m). Scale bar = 0.2 mm. **C**. The same ovule as in Fig. 2b, showing nucellus (n), and integument (i). Scale bar = 0.5 mm. **D**. Detailed view of the ovule in Fig. 2c, showing nucellus (n), micropyle (m), and integument (i). Scale bar = 0.1 mm.

### Remarks

Although with exposed ovules, *Lingyuanfructus* resembles none of reproductive organs in known fossil or living gymnosperms, strobilus or others. This is reinforced by the enclosed ovules seen in the carpels of *Lingyuanfructus* (Figs. 1d-f. 2a).

The term “carpel” is preferred to the term “fruit” since no seed coat expected for a seed is seen in the fossil while integument and micropyle expected an ovule are seen in the fossil of *Lingyuanfructus hibrida* (Figs. 2a-d).

The possibility that ovules of another plant incidentally fall over *Lingyuanfructus*, producing an artifact as if *Lingyuanfructus* had exposed ovules, can be easily eliminated as 1) there are two ovules attached to adjacent carpels while no other isolated ovules are seen around in the specimen. 2) normally, ovules would not fall off until maturing as seeds, but what seen in Figs. 1d and 2a-d are apparently in ovule rather than seed stage. 3) Both of the two exposed ovules of *Lingyuanfructus* are physically connected to the carpels (Figs. 1d, 2a). 4) At least one of the two exposed ovules is physically embedded inside the carpel/carpel. In short, it is impossible for the exposed ovules in *Lingyuanfructus* to be from another plant.

Some of the leaf outlines in Fig. 3a are not physically connected to the main fossil, but these leaves are hard to distinguish morphologically from those physically connected to the main fossil, and their orientation suggests possible connections to the main fossil.

**Figure 3.**
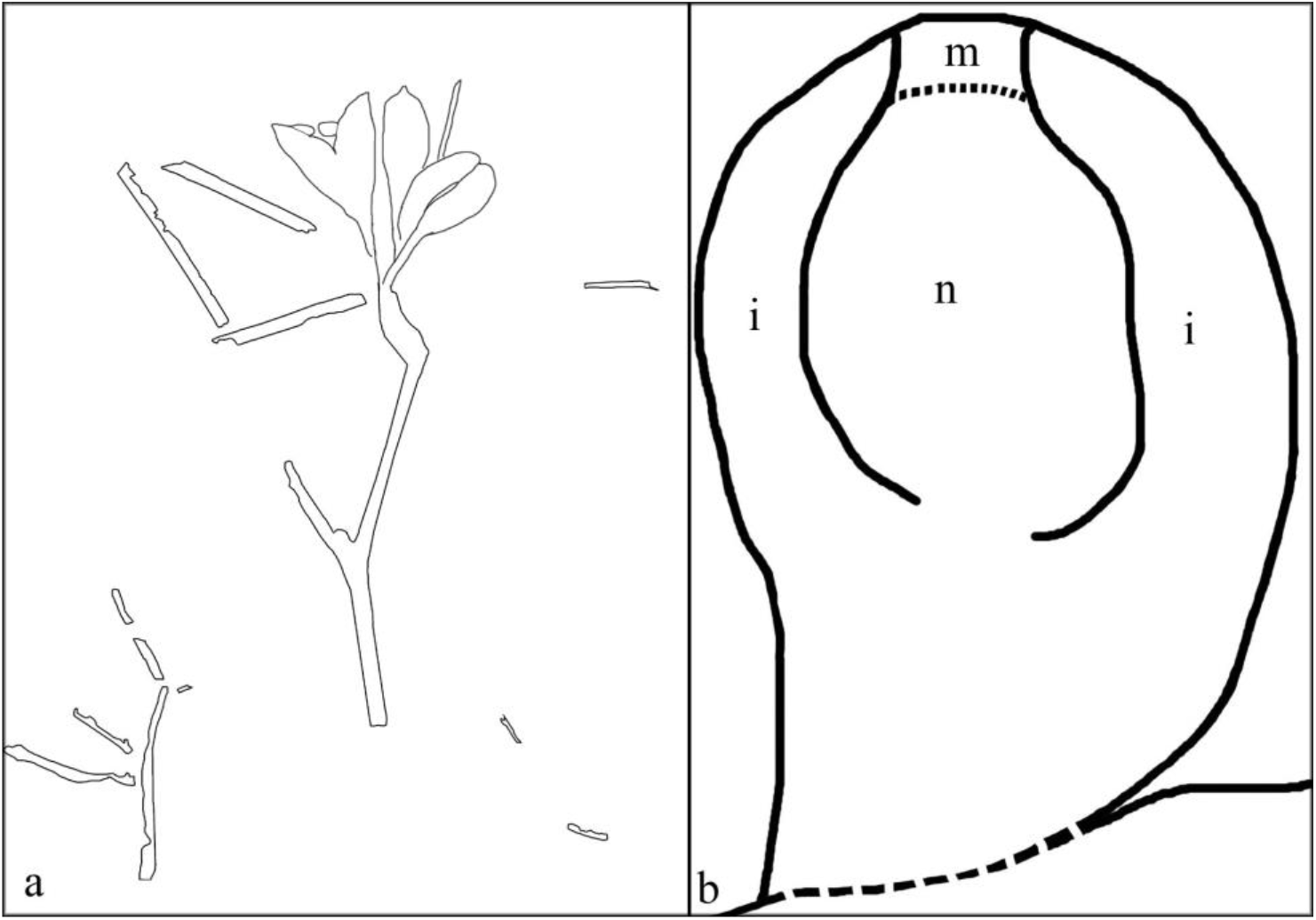
Sketches of *Lingyuanfructus* and its ovule. **A.** Sketch of the specimen shown in Fig. 1a. **B**. Sketch of the exposed orthotropous, unitegmic ovule shown in Fig. 2b that is attached to the carpel margin (bottom), showing nucellus (n), integument (i), and micropyle (m).

### Horizon

the Yixian Formation, Barremian-Aptian, Lower Cretaceous.

### Holotype

PB328298a, PB23898b.

### Etymology

*Lingyuan*-for the fossil locality near Lingyuan, Liaoning, China (41º15ꞌN, 119º15ꞌE); -*fructus* for fruit; *hibrida* for chimeric morphology of the fossil spanning gymnosperms and angiosperms.

### Depository

the Nanjing Institute of Geology and Palaeontology, Nanjing, China.

## Discussions

Literally, angiosperms are characterized by “angiospermy”, which implies that the seeds are enclosed. In this way, the seeds of angiosperms are better protected and nourished, and thus have more chances to give rise to more offspring. A more strict criterion that ensures an angiosperm affinity is angio-ovuly: ovule enclosed before pollination ^8,38^. Since most ovules of *Lingyuanfructus* are enclosed in carpels (Figs. 1d-e, 2a), it is decent to place *Lingyuanfructus* in angiosperms if taking only the enclosed ovules into consideration. However, the intriguing feature of *Lingyuanfructus* is its exposed ovules, which are characteristic of gymnosperms. So it is premature to pin down the final affinity of *Lingyuanfructus* for the time being. In no way, this reduces the systematic significance of *Lingyuanfructus*, instead it boosts the significance of *Lingyuanfructus* in systematics and plant evolution.

Besides enclosed ovules, an intriguing feature of *Lingyuanfructus* is its exposed ovules attached to the adaxial margin of the carpels (Figs. 1a,d-e, 2b-d). The “exposedness” of these ovule implies either 1) that *Lingyuanfructus* is a gymnosperm, or 2) that *Lingyuanfructus*, if an angiosperm, has not fully completed its transition from gymnosperms to angiosperms, and is a step stone between gymnosperms and angiosperms, two otherwise well-separated groups in seed plants.

*Lingyuanfructus*’ orthotropous ovule (Figs. 2b-d) implies that such an ovule is ancestral in angiosperms. This implication is in line with the observation of ovules in the outgroup of angiosperms, gymnosperms. Although bitegmic ovules are frequently seen in angiosperms and their fossil record can be traced back to the Pennsylvanian ^39,40^, ovules in most gymnosperms are unitegmic. The occurrence of unitegmic ovules in *Lingyuanfructus*, of ancestral in angiosperms, is expected as it is suggestive of its proximity to gymnosperms and its distance from crown angiosperms. These two features of *Lingyuanfructus* in tandem point to its position intermediate between gymnosperms and angiosperms.

Although the exposed ovules occur apparently on the adaxial margins of the carpels (Figs. 1d, 2a), the enclosed ovules in *Lingyuanfructus* are attached to both margins of the carpels (Figs. 1d-f). Such placentation has been seen in the model plant *Arabidopsis* (Eudicots) and taken as derived in plant systematics. However, it is noteworthy that such placentation has been seen more than once in early angiosperms. For example, *Neofructus*, another fossil angiosperm from exactly the same fossil locality, also has such placentation ^17^. Not alone, some ovules of *Archaeanthus* (a mid-Cretaceous angiosperm from the Mid-Cretaceous of Kansas, USA) ^41^ are also attached to both margins of the fruits ^42^, not restricted to the adaxial margin. Such observations indicate that the former expectation of placentation for ancestral angiosperms by the traditional theories is spurious and should be updated accordingly.

It is noteworthy that the base and stalk of each carpel in *Lingyuanfructus* are smooth, showing no trace or scar of perianth or stamens (Figs. 1a,d). Such an observation suggests that *Lingyuanfructus* has no typical perianth and no stamens, and thus is pistillate. This underscores the lack of typical flowers in early angiosperms, including *Archaefructus* ^2,3^, *Sinocarpus* ^4,5^, *Baicarpus* ^31^, *Eofructus* ^28^, *Neofructus* ^17^ (all from the Yixian Formation). The consensus of these early angiosperms seems to suggest that these early angiosperms lack true “flowers” typical of living angiosperms. How true “flowers” originated is the next botanical question hard to answer.

## Conclusion

As the first gymno-angiosperm, *Lingyuanfructus* is a step stone between gymnosperms and angiosperms.

## Acknowledgements

I thank Mr. Haisheng Xu for help collecting the specimen, Ms. Jingjing Tang for help with photography using Nikon D800, Ms. Jingyi Yang and Yan Fang for help photography using MAIA3 TESCAN SEM. This research is supported by the Strategic Priority Research Program (B) of Chinese Academy of Sciences (XDB26000000), and National Natural Science Foundation of China (41688103, 91514302, 41572046).

